# Pathogenic and uncertain genetic variants have clinical cardiac correlates in diverse biobank participants

**DOI:** 10.1101/716662

**Authors:** Tess D. Pottinger, Megan J. Puckelwartz, Lorenzo L. Pesce, Avery Robinson, Samuel Kearns, Jennifer A. Pacheco, Laura J. Rasmussen-Torvik, Maureen E. Smith, Rex Chisholm, Elizabeth M. McNally

**Affiliations:** Center for Genetic Medicine, Northwestern University Feinberg School of Medicine, Chicago, Illinois, United States of America; Department of Pharmacology, Northwestern University Feinberg School of Medicine, Chicago, Illinois, United States of America; Computation Institute, University of Chicago, Chicago, Illinois, United States of America; Department of Preventive Medicine, Northwestern University Feinberg School of Medicine, Chicago, Illinois, United States of America; Department of Cell and Molecular Biology, Northwestern University Feinberg School of Medicine, Chicago, Illinois, United States of America

**Keywords:** medically actionable genes, biobank, ClinVar, whole genome sequencing, left ventricle

## Abstract

**Background:** Genome sequencing coupled with electronic heath record data can uncover medically important genetic variation. Interpretation of rare genetic variation and its role in mediating cardiovascular phenotypes is confounded by variants of uncertain significance.

**Methods and Results:** We analyzed the whole genome sequence of 900 racially and ethnically diverse biobank participants selected from a single US center. Participants were equally divided among European, African, Hispanic, and mixed race/ethnicities. We evaluated the American College of Medical Genetics and Genomics medically actionable list of 59 genes focusing on the cardiac genes. Variation was interpreted using the most recent reports in ClinVar, a database of medically relevant human variation. We identified 19 individuals with pathogenic/likely pathogenic variants in cardiac actionable genes (2%) and found evidence for clinical correlates in the electronic health record. African ancestry participants had more variants of uncertain significance in the medically actionable genes including the 30 cardiac actionable genes, even when normalized to total variant count per person. Longitudinal measures of left ventricle size, corrected for body surface area, from approximately 400 biobank participants (1,723 patient years) correlated with genetic findings. The presence of one or more uncertain variants in the actionable cardiac genes and a cardiomyopathy diagnosis correlated with increased left ventricular internal diameter in diastole and in systole. In particular, *MYBPC3* was identified as a gene with excess variants of uncertain significance.

**Conclusions:** These data indicate a subset of uncertain variants may confer risk and should not be considered benign.

## Introduction

Genetic information is increasingly being used in medical decision making, especially for familial cancers and cardiovascular diseases where the identification of rare genetic variants can inform care for patients and family members at risk (1-3). Genetic variants segregating with disease are interpreted as pathogenic or likely pathogenic, and this type of genetic information is diagnostic and useful for clinical management (4). Variants of uncertain significance (VUS) are those genetic variants about which there is insufficient information to adjudicate a pathogenic or benign classification. The VUS designation often arises for rare or unreported missense variants, and this designation is of low medical utility, as its pathogenic status is unknown (4). To improve the reliability of genetic interpretation, the ClinVar database was developed as an online catalog of genetic variation relevant for human health (https://www.ncbi.nlm.nih.gov/clinvar) (5). Genetic testing laboratories regularly contribute to and update ClinVar’s compendium of human health variation. ClinVar, combined with data from large de-identified population sequence databases, is enhancing clinical genetic testing interpretation.

The American College of Medical Genetics and Genomics (ACMG) designated 59 genes as having variation that is medically actionable when the variation is classified as pathogenic or likely pathogenic (6, 7). Not all variation in the ACMG genes is actionable, since some variants are found at high population frequency, thus making their designation benign or likely benign. Uncertain variants are neither pathogenic or benign. However, it can be expected that some of these uncertain variants are, in fact, medically important. It has been recommended to return known pathogenic and likely pathogenic results for the actionable genes, even for biobank participants (8-10). Variants of uncertain significance are not typically reported and returned to biobank participants, as the risks associated with these variants have not been determined. Race and population diversity influence the interpretability of genetic testing results (11-13).

We used whole genome sequencing (WGS) on a diverse cohort of biobank participants from a single metropolitan site in the US. We assessed genetic variation across self-reported race/ethnicity groups, focusing on medically actionable genes and variants previously reported in ClinVar. We found African ancestry participants have a significantly greater number of variants of uncertain significance compared to participants of European descent. This increase in uncertain variants was present across all genes including the ACMG medically actionable genes and the actionable cardiac genes. We now assessed the prevalence of VUS across this diverse population and examined the association of VUS with echocardiographic indicators of cardiac pathology. Uncertain variants in the cardiac medically actionable genes were significantly associated with changes in left ventricular dimensions. These data underscore the complexities of variant interpretation especially in a diverse population.

## Methods

### NUgene Cohort

The NUgene biobank includes adult participants who receive care at Northwestern Medicine. Inclusion and exclusion criteria for participation in the NUgene biobank have been described (14). Through its participation in the eMERGE (Electronic Medical Records and Genomics) consortium (15, 16), the NUgene biobank selected 900 participants for whole genome sequencing (WGS). Race/ethnicity was collected using two methods. Approximately 25% of participants used a questionnaire in which race/ethnicity was selected based on multiple general categories including African, Asian (East and South), European, Hispanic, Pacific Islander, Middle Eastern, and Native American ancestry. Participants who completed this questionnaire could select one or more categories. The remaining 75% of patients were classified based on grandparent ancestry using the general categories named above. As the number of individuals who identified as Asian, Asian Indian, Middle Eastern, and Pacific Islander through both methods were low (<1% of the sample), they were reclassified as “Other” and excluded from further analysis. Individuals who self-reported as multi-racial were classified as mixed, and individuals with at least one grandparent who did not classify similarly to the other three grandparents were classified as mixed. For instance, if an individual selected three grandparents as European descent and one grandparent of African ancestry, this individual is classified as mixed.

### Whole genome sequencing

WGS was performed on an Illumina XTen machine at the Genome Center at Washington University School of Medicine yielding >100 GB of data per sample. This depth correlates with >30-fold coverage across the genome providing more even coverage across both noncoding and coding intervals. WGS data from the NUgene cohort was aligned to the human genome reference sequence GRCh37/hg19 using the Burrows-Wheeler Aligner. Variants were called using the Genome Analysis Tool Kit (GATK v3.3.0) (17, 18). These analyses were conducted using the MegaSeq Pipeline (19).

### Population Structure

The first two components generated by principal component analysis (PCA) were used to estimate global ancestry in the NUgene 900 cohort. PCA was conducted using singular-value decomposition of shared variants in the NUgene cohort with ∼5 million biallelic variants distributed across the genome. Coding and noncoding variants were identified using ANNOVAR (20). Self-reported race was compared to the ancestry groupings determined by the PCA of all variants. An identity by state (IBS) analysis was conducted to identify first relatives within each homogeneous cluster to reduce overfitting in subsequent analytical models. To carry out the IBS analysis, five individuals were removed from the European cluster, one from the African cluster, and four from the Hispanic cluster prior to this analysis to ensure homogeneous race/ethnicity clusters based on visual assessment of PCA graph to classify large outliers. Analyses were carried out using PLINK v1.9 and R v3.5.1.

### ClinVar Analysis

Nonsynonymous coding variants were queried and indexed per genome using ClinVar (February 11^th^, 2019 adjudication of variants from VCF file) (21). These analyses included designations of likely pathogenic, pathogenic, pathogenic/likely pathogenic, uncertain significance, and not reported in ClinVar. The ClinVar VCF file used to annotate these variants only included the most recent designation as determined by February 11, 2019. Generalized regression models adjusting for age and sex were used to determine overall differences as well as pairwise differences between self-reported racial/ethnic groups. To correct for differences in the proportion of variants among the self-reported race/ethnicity groups, variants were normalized based on total variant counts per person (an individual’s variant count was divided by the individual’s total number of variants). Analysis type was determined by the distribution of the outcome variables. In order to correct for multiple testing for the pairwise analysis comparisons, we used the Bonferroni method for correction (22).

### Echocardiogram Analysis with Variants of Uncertain Significance

Echocardiogram, electrocardiogram, and demographic data were queried from the electronic health record, and individual measures were obtained for left ventricular internal diameter-diastole (LVIDd), left ventricular internal diameter-systole (LVIDs), interventricular septal end diastole (IVSDd), and left ventricular ejection fraction (LVEF). The association of longitudinal echocardiogram data and the count of variants of uncertain significance in the cardiac medically actionable genes was calculated. These counts were coded as 0, and ≥1 to reflect the number of variants found in the cardiac actionable genes (as variant counts of 2 were found in only 22 individuals, these counts were recoded as 1). Some participants had multiple echo measurements in a given year. For these participants the median value was used for analysis. A longitudinal model that controlled for age at echo measurement, sex, and self-reported race/ethnicity with an unstructured covariance matrix was used for analysis with the assumption that missing values were missing at random. Year was used as the time component for this analysis. Race/ethnicity specific analyses were also conducted using a similar model. A sensitivity analysis was conducted using individuals who had an ICD-9 code for cardiomyopathy (ICD-9 425 all subcodes) as above. The longitudinal echocardiogram data and the count of variants not identified in the February 2019 ClinVar database, referred to here as unreported, in the cardiac medically actionable genes were also analyzed using a similar method. Analyses were completed using SAS 9.4 and R v3.5.1.

## Results

### Genetic diversity in a US metropolitan health care cohort

Through its participation in the eMERGE (Electronic Medical Records and Genomics) consortium (15, 16), the NUgene biobank selected 900 participants for whole genome sequencing (WGS). The NUgene biobank represents a medical biobank in that the participants receive health care at a single institution. Approximately 23% of these participants were selected based on having diagnostic and/ or procedure codes or medications indicating 1 of 4 conditions of interest to eMERGE investigators: 95 with atopic dermatitis (ICD-9 codes 691.8 and 692.9); 118 with cancer (ICD-9 codes 173-209); 56 with cardiomyopathy (ICD-9 codes 425.1 and 425.4), and 180 with chronic rhinosinusitis (23). The other 77% were selected for race/ethnicity. Whole genome sequencing (WGS) was applied to these 900 diverse individuals, and sequencing reads were aligned to the human genome reference sequence GRCh37/hg19 and variants were called using the MegaSeq Pipeline which utilized BWA and GATK best practices. Of the 900 genomes, five were excluded due to sampling error identified through sex mismatch and/or possible sample contamination. The remaining 895 distributed as follows based on self-reported race/ethnicity: African (26%), European (23%), Hispanic (26%), Mixed (24%), or other (1%) (**Table 1**). Genetic variation correlated with self-reported race/ethnicity. Individuals of European descent had the lowest number of variants per genome compared to those in the other groups (4.9M per European descent genome compared to 5.8M per African descent genome, p< 0.05, **Figure 1A**). Racially mixed and Hispanic descent individuals had average variant counts between African and European ancestry individuals with counts of 5M and 5.4M, respectively. An identity by state (IBS) analysis was performed and, two pairs of first-degree relatives were identified with this analysis.

**Table 1.**
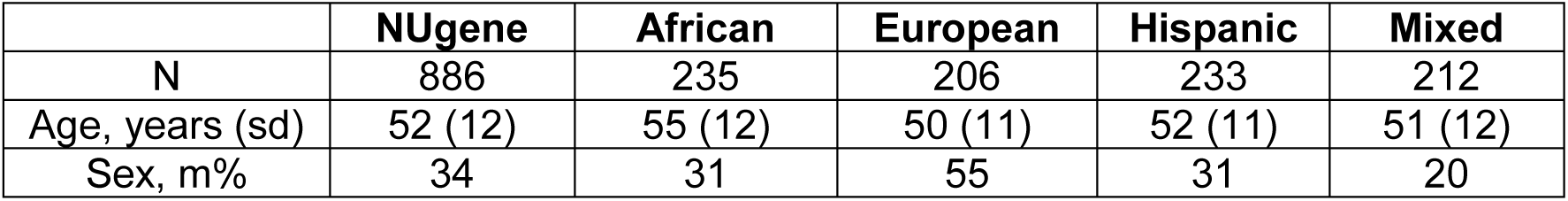
Demographic characteristics of the NUgene Cohort by self-reported race.

**Figure 1.**
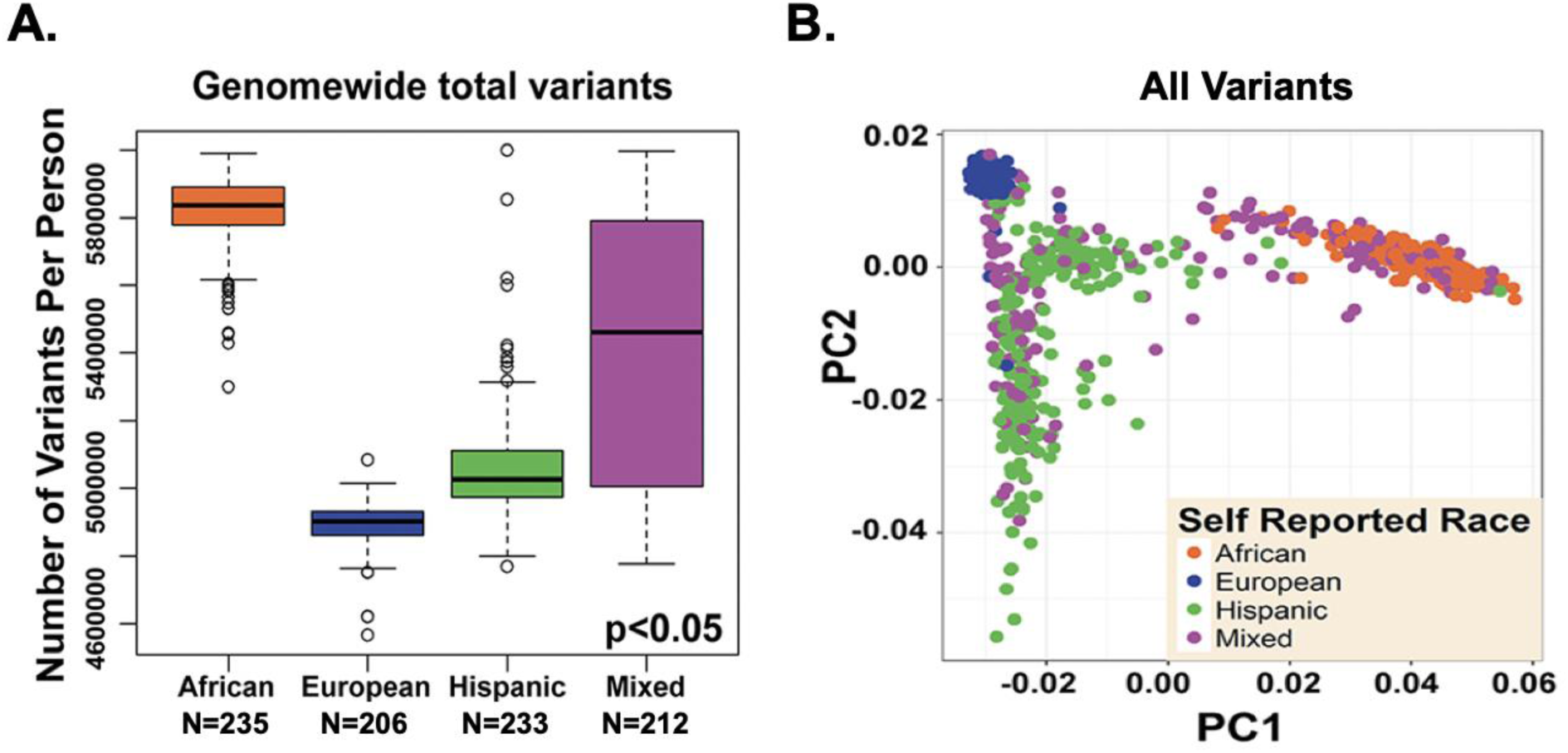
Variant number by race in the NUgene biobank genomes. Consistent with other reports, African ancestry biobank participants had significantly greater genetic variation compared to those of European ancestry. The average number of **(A)** total variants across the genome is shown for each group based on self-reported race; African ancestry (5,815,632±105,545 variants per individual genome); European descent (4,891,014±64,442 variants per individual genome); Hispanic descent (5,056,797±151,424 variants per individual genome) and mixed participants (5,407,116±392,139 variants per individual genome) (p < 0.05, ANOVA across all groups). **(B)** Shown are the results of principal component analysis (PCA) using singular-value decomposition of shared genetic variants among biobank participants. Participants of self-reported African and Hispanic ancestry displayed a more heterogeneous genetic pattern for all variants. Self-reported race (colored circles) are displayed on a genetic clustering background and derived from self-report and grandparent race.

Genomic data was annotated for noncoding and coding variation. As expected, greater than 99% of all genetic variants were noncoding. Considering the nonsynonymous coding variants, the majority were missense variants (79%), and a smaller percentage were stop/loss gain (1.0%), small in-frame insertion/deletions (2.5%), frameshifts (2.5%) and splice sites (15%) (**Supplemental Figure 1**). The number of variants observed only once in the entire data set, a measure of rare variation, was highest in African ancestry participants having, on average, more than any other group (52,865±8,320). European descent participants had the least of any group (30,217±3,992, p<0.05 ANOVA across all groups) (**Supplemental Figure 2**).

**Figure 2.**
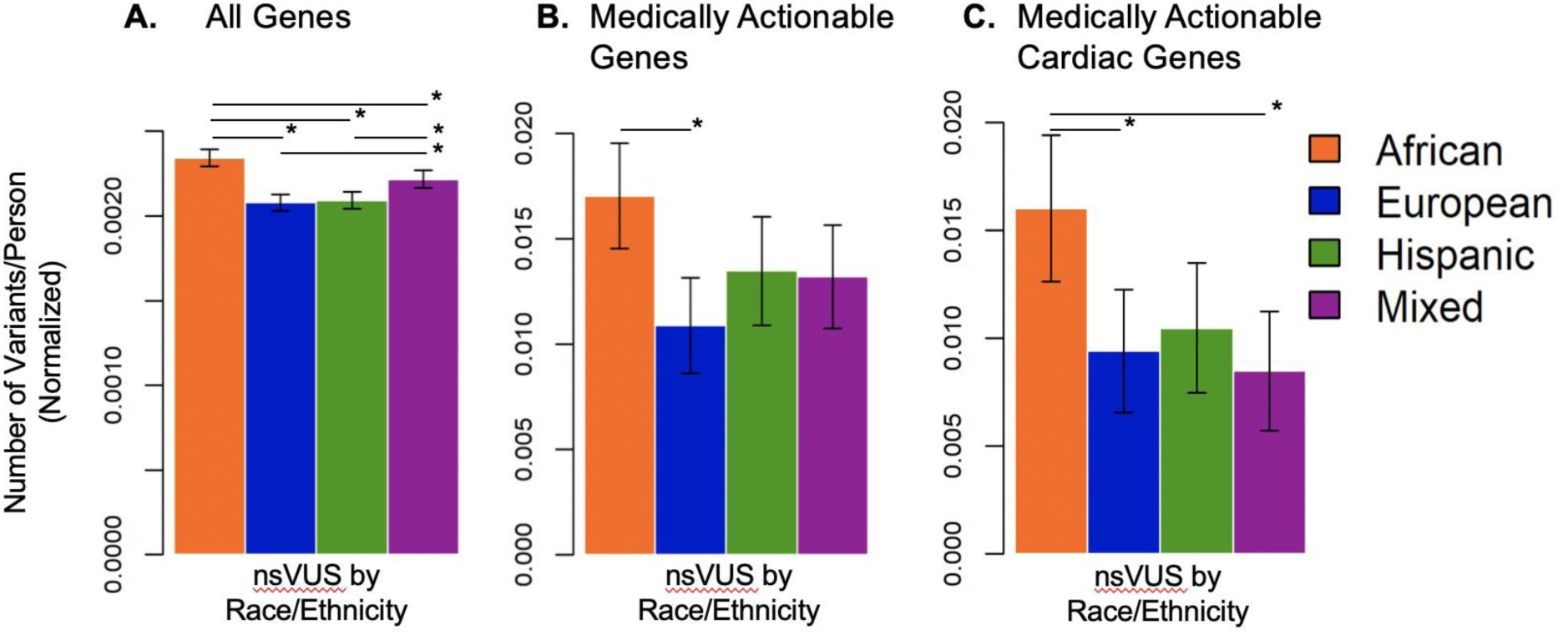
African ancestry biobank participants have significantly more variants of uncertain significance (VUS). **(A)** The average number of total nonsynonymous coding variants across all genes is shown for each group based on self-reported race. Variants of uncertain significance in ClinVar were greater in African ancestry biobank participants compared to other groups (p<0.0001, ANOVA across all groups). **(B)** The average number of nonsynonymous coding variants in the 59 medically actionable genes is shown by self-reported race. The number of variants of uncertain significance reported in ClinVar were greater in African ancestry biobank participants compared to all other groups (p<0.0001, ANOVA across all groups). **(C)** The average number of nonsynonymous coding variants in the 30 cardiac actionable genes is shown by self-reported race. The number of variants of uncertain significance reported in ClinVar were greater in African ancestry biobank participants compared to all other groups (p<0.0001, ANOVA across all groups).

When considering genetic variation across the genome, those who identified as being of European ancestry were tightly clustered on the principal component analysis plot (**Figure 1B**, blue). In contrast, those individuals who identified as African ancestry clustered less tightly (**Figure 1B**, orange). Notably, this group had African and European admixture with as much as 50% European ancestry. Those who identifying as Hispanic had a non-uniform structure with two major groupings, including one that clusters along the Euro-Asian line, and the other without a clear racial grouping indicating a more mixed ancestry (**Figure 1B**, green).

### African ancestry subjects have more variants of uncertain significance

Genetic testing is increasingly being used in adult health care settings, but the interpretation of genetic results is complicated by the rare frequency of many genetic variants (24, 25). Genetic testing is further complicated by the observations that pathogenic and likely pathogenic variants are found at higher frequencies than the diseases specified by these variants (26, 27). We queried the number of nonsynonymous variants in the NUgene biobank genomes that were previously reported in ClinVar, a database of clinically relevant genetic information (5). The most recent variant adjudication was used for this analysis; approximately 90% of variants were adjudicated since 2015, the time when the ACMG guidelines were released (6). African ancestry individuals are known to have greater genetic variation than either their European or Hispanic counterparts. Therefore, variant counts were divided by total variant count per person to account for this baseline difference across populations (28). NUgene participants of African ancestry had more VUS compared to individuals of European, Hispanic and mixed ancestry even after normalization (p < 0.0001, **Figure 2A**).

The NUgene genomes were then queried for nonsynonymous variation in the 59 medically-actionable genes (**Figure 2B**) (7). The mean normalized number of pathogenic and likely pathogenic variants in the 59 medically actionable gene lists did not differ across groups; however, the total number of these variants was very small. Within the medically-actionable genes, NUgene individuals of African ancestry were more likely to have VUS than individuals of European ancestry even after normalization to total variant count per person (p < 0.05, **Figure 2B**). VUS numbers were similar between the other ancestry groups (**Figure 2B**). The medically actionable genes were subdivided into cardiac and cancer genes and similarly analyzed (**Supplemental Table 1**). For cardiac medically actionable genes, African ancestry participants had more VUSs compared to European and mixed individuals even after normalization to total variant count (p < 0.05) (**Figure 2C**).

### Medically actionable cardiac genes variants of uncertain significance correlate with left ventricular measures

Of the medically actionable genes, 30 are linked to cardiovascular conditions (**Supplemental Table 1**). The number of participants with ClinVar adjudicated pathogenic and likely pathogenic (P/LP) variants in these genes totaled 19 with no single individual having more than one P/LP variant (**Table 2**). Of these 19 individuals, 13 had an electrocardiogram, echocardiogram or test of their cholesterol level in the electronic health record, and those with electronic health cardiac data were ten years older than those without electronic health cardiac information (52 vs 42 years). Of the four individuals who had P/LP variants in long QT syndrome genes, all had a prolonged QTc (**Table 3**).

**Table 2.** Genetic Variation in the NUgene Cohort derived classified by ClinVar.

| **Pathogenic Variants** | **Total** | **Mean/Person** | **Range** |
| --- | --- | --- | --- |
| All genes | 886 | 6.51 | 1 - 19 |
| Medically actionable genes | 23 | 0.026 | 0 - 1 |
| Cardiac actionable genes | 10 | 0.011 | 0 - 1 |
| **Likely Pathogenic Variants** | **Total** | **Mean/Person** | **Range** |
| All genes | 660 | 1.02 | 0 - 4 |
| Medically actionable genes | 7 | 0.0079 | 0 - 1 |
| Cardiac actionable genes | 4 | 0.0045 | 0 - 1 |
| **Variants of Uncertain Significance** | **Total** | **Mean/Person** | **Range** |
| All genes | 886 | 33.42 | 15 - 58 |
| Medically actionable genes | 385 | 0.58 | 0 - 4 |
| Cardiac actionable genes | 191 | 0.24 | 0 - 3 |
| **Unreported Variants** | **Total** | **Mean/Person** | **Range** |
| All genes | 886 | 13,430 | 10,042 - 15,282 |
| Medically actionable genes | 886 | 2.57 | 2 - 13 |
| Cardiac actionable genes | 886 | 2.24 | 2 - 6 |

**Table 3.**
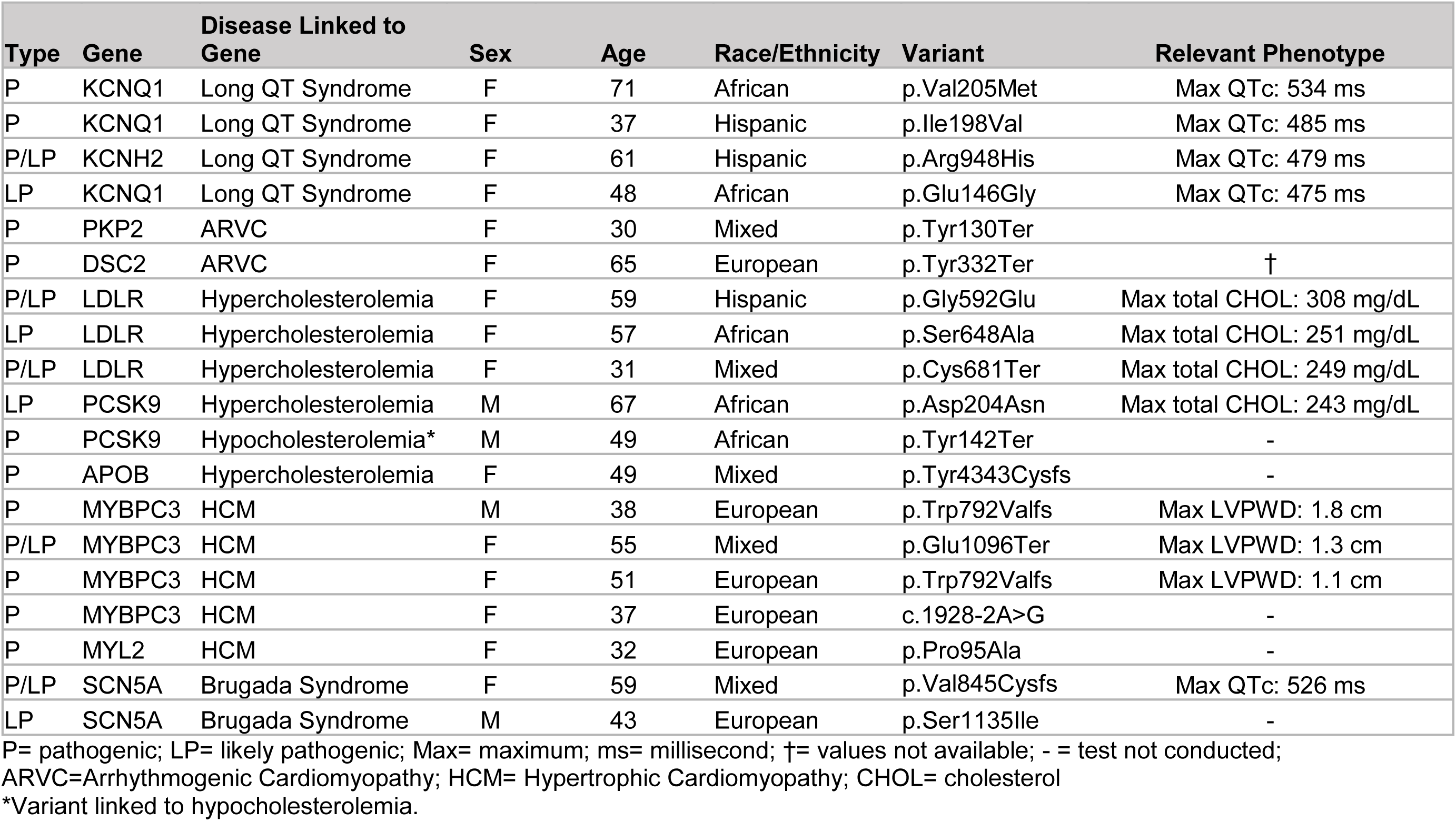
Pathogenic and Likely Pathogenic variants found in Cardiac Actionable Genes in the NUgene Cohort and available electronic health record data. Table 4. Longitudinal Association of Left Ventricular Measures with VUS in Cardiac Actionable Genes from 94 Subjects with Cardiomyopathy Diagnostic Codes SIGNIFICANT VALUES *** p<0.001, ** p<0.01, * p<0.05;

| Type | Gene | Disease Linked to Gene | Sex | Age | Race/Ethnicity | Variant | Relevant Phenotype |
| --- | --- | --- | --- | --- | --- | --- | --- |
| P | KCNQ1 | Long QT Syndrome | F | 71 | African | p.Val205Met | Max QTc: 534 ms |
| P | KCNQ1 | Long QT Syndrome | F | 37 | Hispanic | p.Ile198Val | Max QTc: 485 ms |
| P/LP | KCNH2 | Long QT Syndrome | F | 61 | Hispanic | p.Arg948His | Max QTc: 479 ms |
| LP | KCNQ1 | Long QT Syndrome | F | 48 | African | p.Glu146Gly | Max QTc: 475 ms |
| P | PKP2 | ARVC | F | 30 | Mixed | p.Tyr130Ter |  |
| P | DSC2 | ARVC | F | 65 | European | p.Tyr332Ter | † |
| P/LP | LDLR | Hypercholesterolemia | F | 59 | Hispanic | p.Gly592Glu | Max total CHOL: 308 mg/dL |
| LP | LDLR | Hypercholesterolemia | F | 57 | African | p.Ser648Ala | Max total CHOL: 251 mg/dL |
| P/LP | LDLR | Hypercholesterolemia | F | 31 | Mixed | p.Cys681Ter | Max total CHOL: 249 mg/dL |
| LP | PCSK9 | Hypercholesterolemia | M | 67 | African | p.Asp204Asn | Max total CHOL: 243 mg/dL |
| P | PCSK9 | Hypocholesterolemia* | M | 49 | African | p.Tyr142Ter | - |
| P | APOB | Hypercholesterolemia | F | 49 | Mixed | p.Tyr4343Cysfs | - |
| P | MYBPC3 | HCM | M | 38 | European | p.Trp792Valfs | Max LVPWD: 1.8 cm |
| P/LP | MYBPC3 | HCM | F | 55 | Mixed | p.Glu1096Ter | Max LVPWD: 1.3 cm |
| P | MYBPC3 | HCM | F | 51 | European | p.Trp792Valfs | Max LVPWD: 1.1 cm |
| P | MYBPC3 | HCM | F | 37 | European | c.1928-2A>G | - |
| P | MYL2 | HCM | F | 32 | European | p.Pro95Ala | - |
| P/LP | SCN5A | Brugada Syndrome | F | 59 | Mixed | p.Val845Cysfs | Max QTc: 526 ms |
| LP | SCN5A | Brugada Syndrome | M | 43 | European | p.Ser1135Ile | - |

P= pathogenic; LP= likely pathogenic; Max= maximum; ms= millisecond; †= values not available; - = test not conducted;
ARVC=Arrhythmogenic Cardiomyopathy; HCM= Hypertrophic Cardiomyopathy; CHOL= cholesterol
\*Variant linked to hypocholesterolemia.

**Table 4.** Longitudinal Association of Left Ventricular Measures with VUS in Cardiac Actionable Genes from 94 Subjects with Cardiomyopathy Diagnostic Codes SIGNIFICANT VALUES *** p<0.001, ** p<0.01, * p<0.05;

| Echo Measure<br>VUS ≥1 vs<br>VUS=0 | Total<br>Observations | African | European | Hispanic | Mixed |
| --- | --- | --- | --- | --- | --- |
| **LVIDd/BSA** | n=447 | n=99 | n=238 | n=42 | n=68 |
| Slope Difference<br>Estimate | **+0.035 \*\*** | +0.0012 | **+0.038 \*** | **+0.015 \*\*\*** | - |
| **LVIDs/BSA** | n=442 | n=94 | n=239 | n=41 | n=68 |
| Slope Difference<br>Estimate | **+0.047 \*\*\*** | **+0.047 \*\*** | **+0.05 \*** | - | -0.0035 |
| **LVEF** | n=426 | n=90 | n=225 | n=41 | n=70 |
| Slope Difference<br>Estimate | -0.42 | **-1.93 \*\*** | -0.25 | - | +2.19 |
| **IVSDd/BSA** | n=465 | n=101 | n=248 | n=45 | n=71 |
| Slope Difference<br>Estimate | -0.0014 | -0.013 † | +0.008 | -0.016 | +0.0065 |

† p<0.1
Slope difference estimates compared ≥1 VUS to 0 VUS, and the model controlled for age at echocardiogram, sex, and self-reported race/ethnicity
LVIDd=left ventricular diameter diastole; BSA=body surface area; IVSDd=interventricular septal end diastole
LVIDs=left ventricular diameter systole; LVEF=left ventricular ejection fraction

Because a proportion of VUS are likely to confer phenotype, VUS within the medically-actionable cardiac genes were analyzed for their association with echocardiographic measures. Of the 385 individuals with echocardiographic data in the electronic health records, 191 individuals had at least one VUS in the cardiac genes (**Table 2**). Of these cardiac actionable genes, *MYBPC3* had the most VUSs (**Figure 3**). To determine if VUS count was associated with cardiac phenotype measures across the cohort, a Loess plot was used to compare left ventricular measures over time, corrected for body surface area (BSA), and then correlated with presence of VUSs. Longitudinal measures of left ventricular internal diameter in diastole (LVIDd) and left ventricular internal diameter in systole (LVIDs) increased with VUS count (**Supplemental Figure 3**). Longitudinal analyses revealed a significant increase in LVIDd over time in those with ≥1 VUS compared to those without VUSs in cardiac genes, and this significance was observed after controlling for age at echocardiogram, sex and self-reported race/ethnicity (p<0.05, **Supplemental Table 2**). When evaluating this same data within each race/ethnic group, a similar trend was seen for LVIDd and VUS count for participants of European and Hispanic descent. Individuals with cardiac VUSs also showed a similar effect for systolic measurements (LVIDs) over time (p<0.01, **Supplemental Table 2**). In this case, the race/ethnic specific analysis identified those of African and European ancestry as having a significant change over time (p<0.05). Left ventricular ejection fraction (LVEF) and interventricular septal end diameter-diastole (IVSDd) showed no significant differences in the pooled analyses. However, in the race/ethnicity analysis, African ancestry individuals with VUSs had a significant decrease in LVEF over time (p<0.05, **Supplemental Table 2**). These data indicate that the presence of VUSs within cardiac actionable genes associates with a change in cardiac dimensions over time.

**Figure 3.**
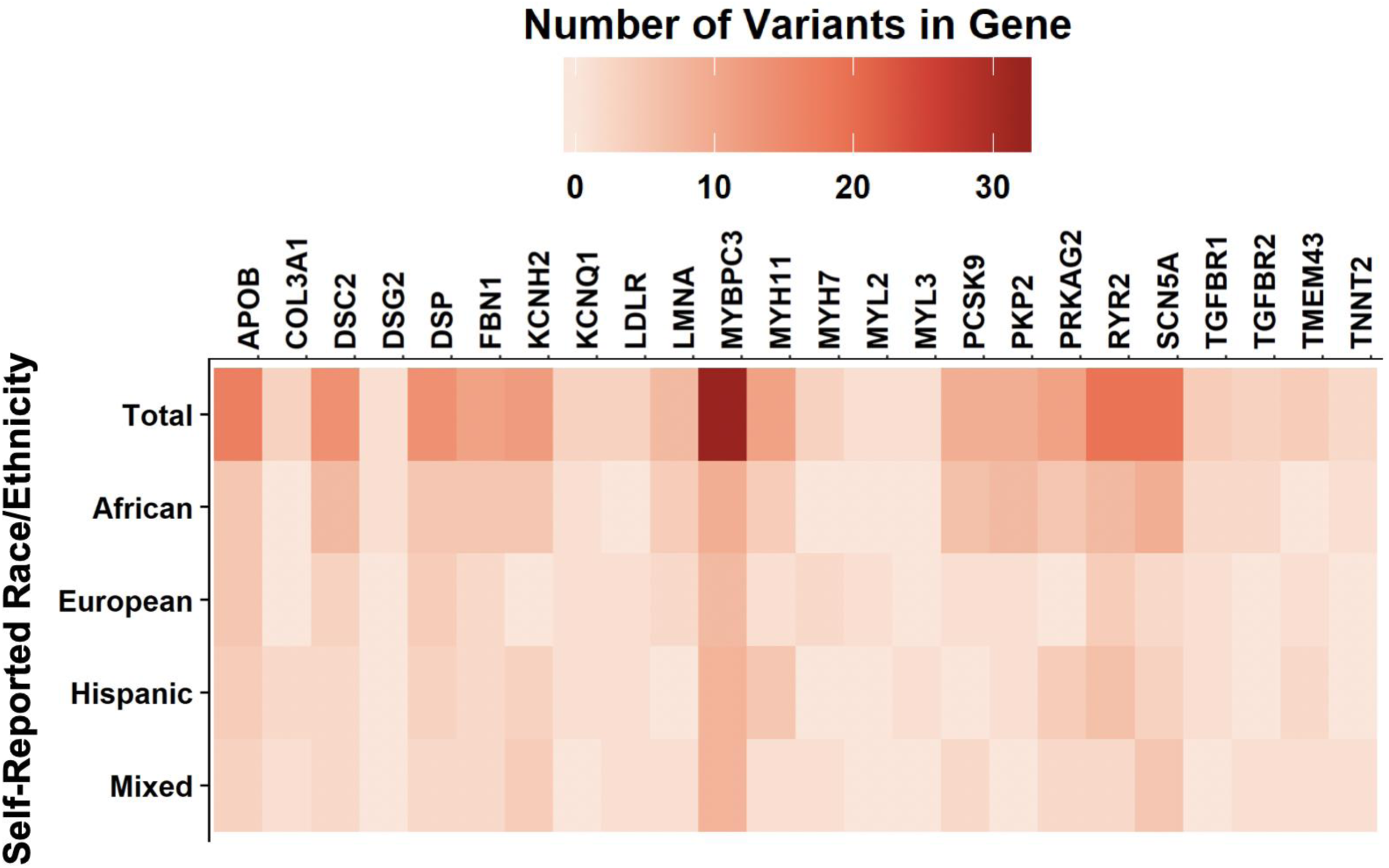
Number of Variants of Uncertain Significance (VUS) in medically actionable cardiac genes. VUS number in the cardiac actionable genes is indicated across the NUgene Cohort (total cohort in top line), and by self-reported race/ethnicity in each line below.

To determine if these trends were driven by subjects with diagnosed cardiomyopathy, the electronic health record was queried for ICD-9 codes for cardiomyopathy including ICD-9 425. Ninety-four subjects were identified, including the original 56 pre-selected at the time of sequencing. The change in LV dimensions over time observed for the 94 cardiomyopathy subjects appeared similar to that of the entire cohort (**Figure 4** and **Table 4**). However, none of these 94 individuals had a known pathogenic variant in the cardiac medically actionable gene list. Because there are more cardiomyopathy genes beyond those on the medically actionable list, we queried variation in 102 cardiomyopathy genes; this list of cardiomyopathy genes was derived from gene panels used in commercial testing labs (**Supplemental Table 4**). Only one individual had a pathogenic variant in the cardiac panel, *TTR* V122I, and a VUS in the cardiac medically actionable genes. Additional subjects with pathogenic variants lacked VUSs in the cardiac medically actionable genes and, therefore did not contribute to changes in LV dimensions seen in **Figure 4**. Twenty-four cardiomyopathy-diagnosed individuals harbored VUSs in cardiomyopathy genes, and these individuals are responsible for the change in LV dimensions over time seen in **Figure 4**. The list of these VUSs is shown in **Supplemental Table 5 and 6**. Five of 24 had VUSs in *MYBPC3*. These data suggest that VUSs contribute to cardiomyopathy and highlight the importance of interpreting variant pathogenicity in the context of phenotype.

**Figure 4.**
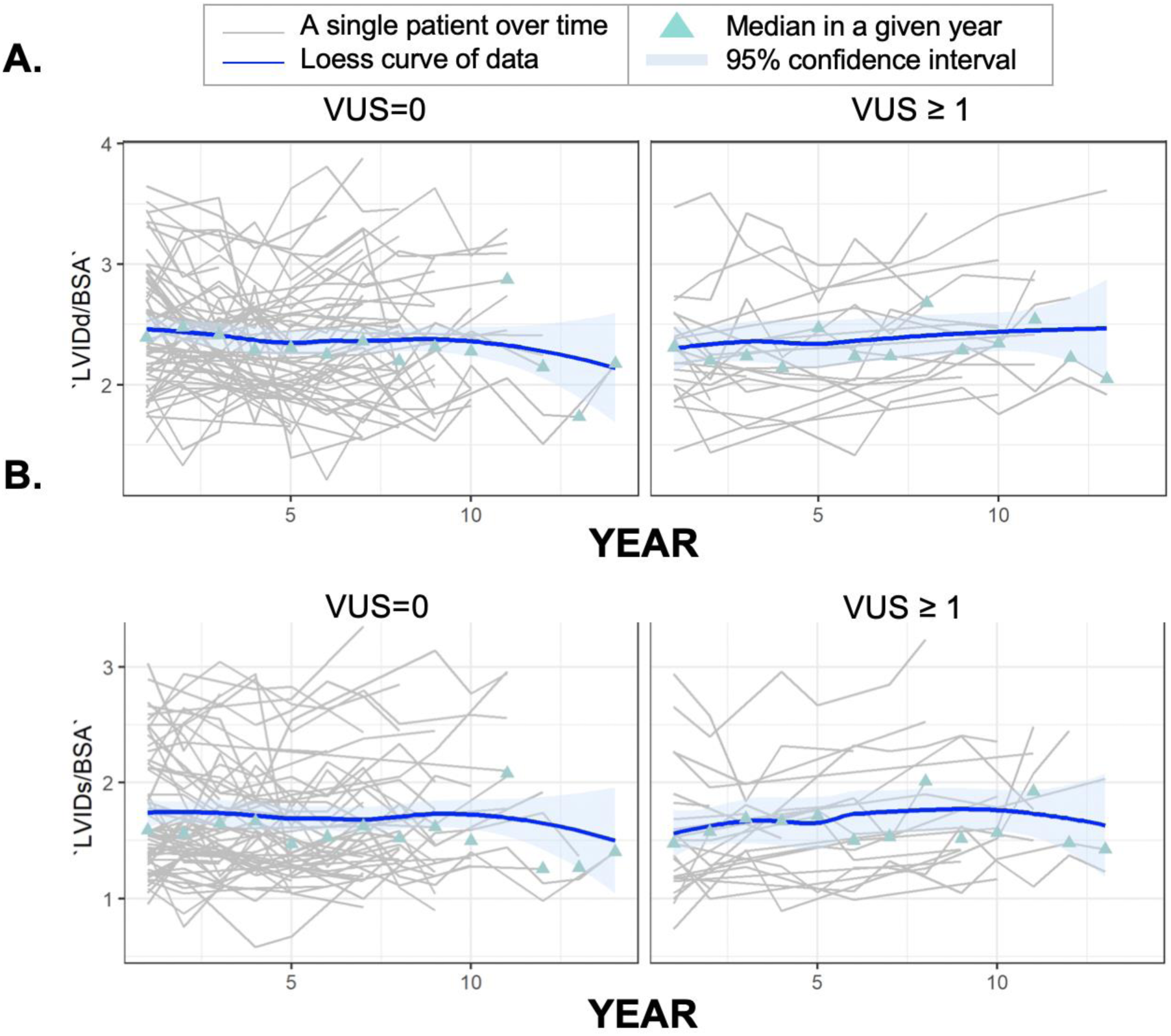
Median left ventricular dimensions correlated with VUS number in the cardiac actionable genes in 94 patients with a cardiomyopathy diagnosis. **(A)** Left ventricular internal diameter in diastole (LVIDd) corrected for body surface area (BSA) over 14 years of hospital visits by the number of variants of uncertain significance found in the cardiac actionable genes. **(B)** Left ventricular internal diameter in systole (LVIDs) corrected for BSA over 14 years of hospital visits by the number of variants of uncertain significance found in the cardiac actionable genes.

### Evaluating variants unreported in ClinVar from diverse biobank participants

We next examined variants not previously reported in ClinVar. Across all genes, participants of African ancestry had more variants not previously reported in ClinVar (p < 0.0001, **Supplemental Figure 4A**). For the medically actionable genes, there were similar numbers of variants not previously reported in ClinVar among all groups, when variant counts were normalized to the total number of variants per person found in these genes (**Supplemental Figure 4B**). For cardiac actionable genes, African ancestry participants had fewer unreported variants compared to individuals of European and Hispanic ancestry when normalized to the total number of variants found in these genes (p < 0.0001 for both) (**Supplemental Figure 4C**).

We evaluated the relationship between echocardiographic measurements and unreported variants in the cardiac medically-actionable genes (**Table 2**). The genes with the most unreported variants were *APOB, MYH11*, and *DSP* (**Supplemental Figure 5**). Longitudinal analyses of LVIDd, LVIDs, IVSDd, and LVEF, corrected for BSA, showed no difference over time when evaluating unreported rare variants. These analyses controlled for age at echocardiogram, sex, and self-reported race/ethnicity (**Supplemental Table 3**). While there may be variants of clinical impact in this dataset, the signal may be masked by the large number of benign variants.

## Discussion

### Clinically actionable findings in diverse biobank participants

The utility of genetic information improves with deeper and more diverse genetic databases. This principle underlies All-of-Us, and the Million Veteran programs, which aim to provide a broader genetic picture of the diverse US population. For hypertrophic cardiomyopathy, it was previously suggested that the under-representation of participants from diverse racial and ethnic backgrounds in large public databases led to the misclassification of benign variants as pathogenic, especially in non-European ancestry populations (26). In this study, individuals with pathogenic or likely pathogenic variants in the cardiac actionable genes had evidence of cardiac clinical findings in the electronic health record, and this was exemplified by the four diverse biobank participants with variants in genes linked to long QT syndrome. Because these variants increase risk for sudden death (29), these genetic findings represent an opportunity for risk reduction. These findings differ from a previous report where rare long QT syndrome-associated variants did not display clear differences from controls (30). However, this current analysis relied more on ClinVar for interpretation, and during the intervening three years between the prior study and now, ClinVar has expanded with data contributions from more clinical testing and now includes many more variants interpreted through more consistent guidelines.

### Racial differences in unreported variants

Given its composition, ClinVar’s catalog of genetic variation, in part, reflects genetic testing practices (5). The findings that European descent individuals are less likely to have unreported variants than other race/ethnic groups (p<0.001) may reflect that clinical genetic testing is disproportionally applied to this group. But this observation also likely reflects the smaller degree of genetic diversity within this group. Genetic diversity may contribute to disparity in interpreting genetic testing results in non-European and especially African ancestry individuals (31, 32). This study showed that variants of uncertain significance were disproportionally higher in individuals of African ancestry than all other individuals, and this trend continued when the cardiac medical actionable genes were analyzed. However, this study also showed that individuals of African ancestry had fewer unreported variants in cardiac medically actionable genes when normalized to the total number of variants per person found in these genes. This may indicate that this disparity is more significant for genes which are not currently considered medically actionable. In addition, this information could also reflect the complexity in adjudicating variants in individuals of non-European ancestries, as this process is complicated by the high number of rare or private variants found in these populations, which is well positioned to contribute to interpretation as a VUS.

### Variants of uncertain significance and echocardiographic findings

This study also identified echocardiographic findings associated with the presence of VUSs, suggesting that some VUSs influence cardiac phenotype. Left ventricular internal diameters, in diastole and systole, correlated with having ≥1 VUSs, and this result was especially evident in those with a cardiomyopathy diagnosis in the electronic health record. An array of cardiomyopathy genes, and other cardiac genes, harbored VUSs in these subjects, and *MYBPC3* was noted as having the highest number of VUSs. *MYBPC3* truncations are a common cause of hypertrophic cardiomyopathy (1, 33), and the *MYBPC3* VUSs were all missense, perhaps suggesting a distinct mode of action. In context of genetic testing for cardiomyopathies, variants of uncertain significance are typically returned and can sometimes be interpreted with further familial testing and segregation analysis (34, 35). In the setting of biobank testing, such results would not be returned to subjects or providers, in part because interpretation of pathogenic variation is done outside the context of phenotype. We observed the correlation of VUS with LV dimensions, when viewing the entire cohort, but this finding was evident when considering only those with a cardiomyopathy diagnosis. The identification of genetic variants contributing to cardiomyopathy affects management, especially for the accompanying arrhythmia risk (1). The inability to fully interpret these variants limits the use of this data for both the patients and their family members. Improved methods in which variants are interpreted in concert with clinical diagnoses may address this deficiency.

Uptick of genetic testing in the diverse clinical setting, along with stricter guidelines on interpretation, and the limitations of *in silico* tools have contributed to an increased number of uncertain variants (36, 37). In practice, the enrichment of VUS in specific racial groups makes genetic testing harder to interpret within those groups. Expanding genetic databases to include self-reported race and ultimately linking this data to health information will facilitate genetic interpretation. The dataset developed in this report, along with the additional data generated from the eMERGE consortium, extends the diversity of publicly-available genetic information (16). Until databases are sufficiently powered to address these deficiencies, in some cases, it may be reasonable to return VUS results to biobank participants, provided the biobank participants wish to receive these results. This would allow participants and their healthcare providers to assess risk by integrating this genetic data with personal medical findings and family medical history.

## Acknowledgements

We thank the McDonnell Genome Institute at Washington University in St. Louis. We also gratefully acknowledge the participation of the NUgene biobank participants.

## Supporting information

**Supplemental Table 1.** Medically Actionable Genes by Cardiac and Cancer Designation.

| Cardiac Genes | Cancer Genes | Other Genes |
| --- | --- | --- |
| *ACTA2* | *APC* | *ATP7B* |
| *ACTC1* | *BMPR1A* | *CACNA1S* |
| *APOB* | *BRCA1* | *OTC* |
| *COL3A1* | *BRCA2* | *RYR1* |
| *DSC2* | *MEN1* |  |
| *DSG2* | *MLH1* |  |
| *DSP* | *MSH2* |  |
| *FBN1* | *MSH6* |  |
| *GLA* | *MUTYH* |  |
| *KCNH2* | *NF2* |  |
| *KCNQ1* | *PMS2* |  |
| *LDLR* | *PTEN* |  |
| *LMNA* | *RB1* |  |
| *MYBPC3* | *RET* |  |
| *MYH11* | *SDHAF2* |  |
| *MYH7* | *SDHB* |  |
| *MYL2* | *SDHC* |  |
| *MYL3* | *SDHD* |  |
| *PCSK9* | *SMAD4* |  |
| *PKP2* | *STK11* |  |
| *PRKAG2* | *TP53* |  |
| *RYR2* | *TSC1* |  |
| *SCN5A* | *TSC2* |  |
| *SMAD3* | *VHL* |  |
| *TGFBR1* | *WT1* |  |
| *TGFBR2* |  |  |
| *TMEM43* |  |  |
| *TNNI3* |  |  |
| *TNNT2* |  |  |
| *TPM1* |  |  |

**Supplemental Table 2.** Longitudinal Association of Left Ventricular Measures with VUS number in the Cardiac Actionable Genes <u>in the entire cohort</u>.SIGNIFICANT VALUES *p<0.05, **p<0.01.

| **Echo Measure**<br>**VUS ≥1 vs VUS=0** | **Total**<br>**Observations** | **African** | **European** | **Hispanic** | **Mixed** |
| --- | --- | --- | --- | --- | --- |
| **LVIDd/BSA** | n=914 | n=266 | n=316 | n=151 | n=181 |
| Slope Difference Estimate | **+0.018 \*** | +0.0021 | **+0.034 \*** | +0.042† | +0.025 |
| **LVIDs/BSA** | n=984 | n=277 | n=327 | n=169 | n=211 |
| Slope Difference Estimate | **+0.027 \*\*** | **+0.026 \*** | **+0.047 \*** | +0.027 | +0.0058 |
| **LVEF** | n=987 | n=273 | n=319 | n=166 | n=229 |
| Slope Difference Estimate | -0.33 | **-0.85 \*** | -0.32 | +0.036 | +0.36 |
| **IVSDd/BSA** | n=1,049 | n=296 | n=341 | n=185 | n=227 |
| Slope Difference Estimate | -0.00068 | -0.0063 | +0.0092† | -0.0063 | -0.0055 |

† p<0.1
Slope difference estimates compared 1 or more VUS to 0 VUS, and the model controlled for age at echocardiogram, sex, and self-reported race/ethnicity.
LVIDd=left ventricular diameter diastole; BSA=body surface area; IVSDd=interventricular septal end diastole
LVIDs=left ventricular diameter systole; LVEF=left ventricular ejection fraction

**Supplemental Table 3.** Longitudinal Association of Left Ventricular Measures with <u>Unreported Variants</u> in Cardiac Actionable Genes <u>in the entire cohort</u>.

| **Echo Measure**<br>**VUS ≥1 vs**<br>**VUS=0** | **Total**<br>**Observations** | **African** | **European** | **Hispanic** | **Mixed** |
| --- | --- | --- | --- | --- | --- |
| **LVIDd/BSA** | n=914 | n=266 | n=316 | n=151 | n=181 |
| Slope Difference Estimate | -0.069 | -0.016 | +0.0035 | -0.028 | +0.012 |
| **LVIDs/BSA** | n=984 | n=277 | n=327 | n=169 | n=211 |
| Slope Difference Estimate | +0.0058 | +0.012 | +0.019 | -0.028 | -0.0036 |
| **LVEF** | n=987 | n=273 | n=319 | n=166 | n=229 |
| Slope Difference Estimate | +0.35 | +0.8 † | -0.3 | +0.29 | +0.47 |
| **IVSDd/BSA** | n=1,049 | n=296 | n=341 | n=185 | n=227 |
| Slope Difference Estimate | -0.0013 | +0.0043 | -0.0074 | +0.0074 | -0.0072 |

**Supplemental Table 4.**
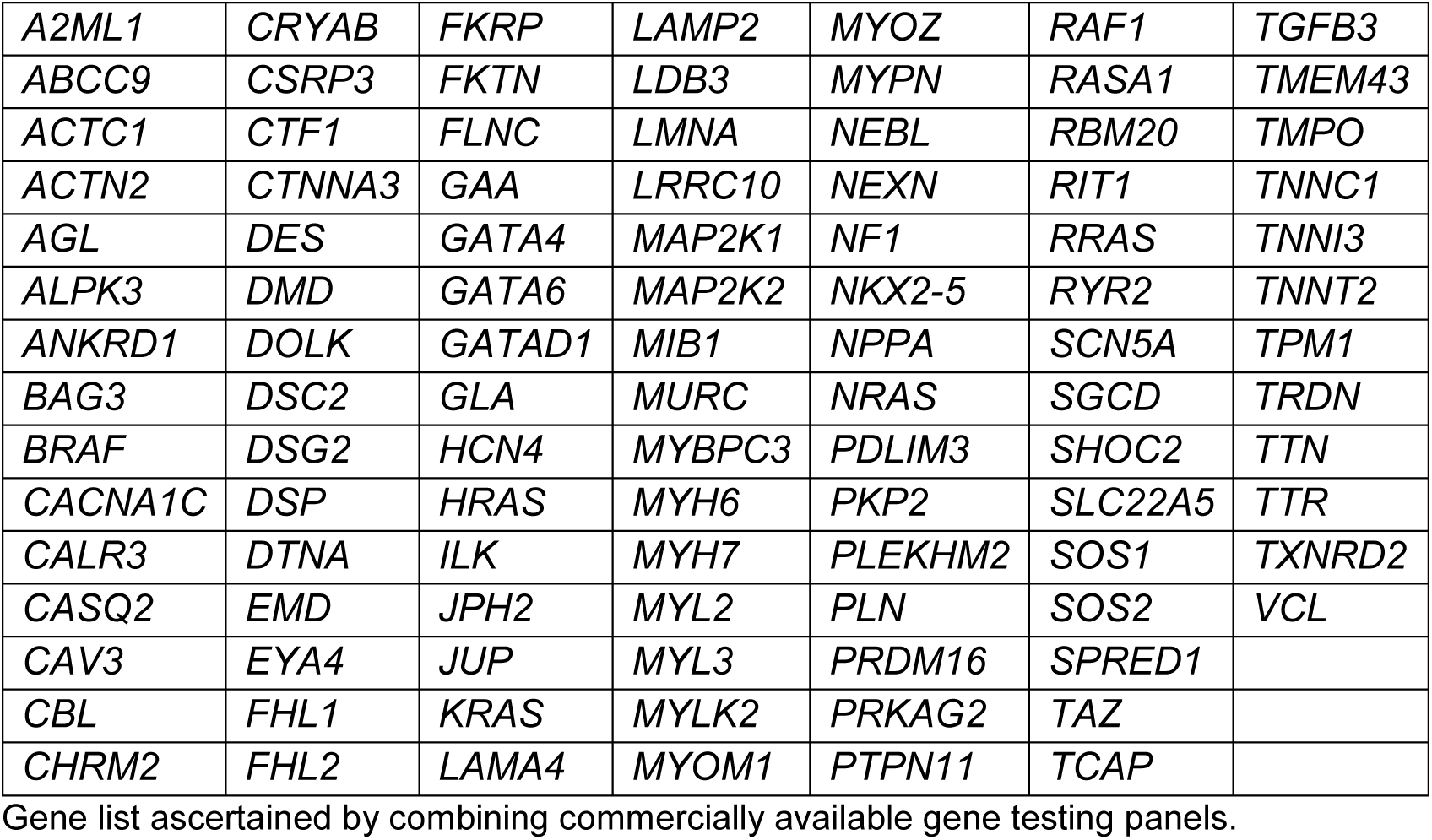
Cardiomyopathy Genes.

|  |  |  |  |  |  |  |
| --- | --- | --- | --- | --- | --- | --- |
| *A2ML1* | *CRYAB* | *FKRP* | *LAMP2* | *MYOZ* | *RAF1* | *TGFB3* |
| *ABCC9* | *CSRP3* | *FKTN* | *LDB3* | *MYPN* | *RASA1* | *TMEM43* |
| *ACTC1* | *CTF1* | *FLNC* | *LMNA* | *NEBL* | *RBM20* | *TMPO* |
| *ACTN2* | *CTNNA3* | *GAA* | *LRRC10* | *NEXN* | *RIT1* | *TNNC1* |
| *AGL* | *DES* | *GATA4* | *MAP2K1* | *NF1* | *RRAS* | *TNNI3* |
| *ALPK3* | *DMD* | *GATA6* | *MAP2K2* | *NKX2-5* | *RYR2* | *TNNT2* |
| *ANKRD1* | *DOLK* | *GATAD1* | *MIB1* | *NPPA* | *SCN5A* | *TPM1* |
| *BAG3* | *DSC2* | *GLA* | *MURC* | *NRAS* | *SGCD* | *TRDN* |
| *BRAF* | *DSG2* | *HCN4* | *MYBPC3* | *PDLIM3* | *SHOC2* | *TTN* |
| *CACNA1C* | *DSP* | *HRAS* | *MYH6* | *PKP2* | *SLC22A5* | *TTR* |
| *CALR3* | *DTNA* | *ILK* | *MYH7* | *PLEKHM2* | *SOS1* | *TXNRD2* |
| *CASQ2* | *EMD* | *JPH2* | *MYL2* | *PLN* | *SOS2* | *VCL* |
| *CAV3* | *EYA4* | *JUP* | *MYL3* | *PRDM16* | *SPRED1* |  |
| *CBL* | *FHL1* | *KRAS* | *MYLK2* | *PRKAG2* | *TAZ* |  |
| *CHRM2* | *FHL2* | *LAMA4* | *MYOM1* | *PTPN11* | *TCAP* |  |

Gene list ascertained by combining commercially available gene testing panels.

**Supplemental Table 5.**
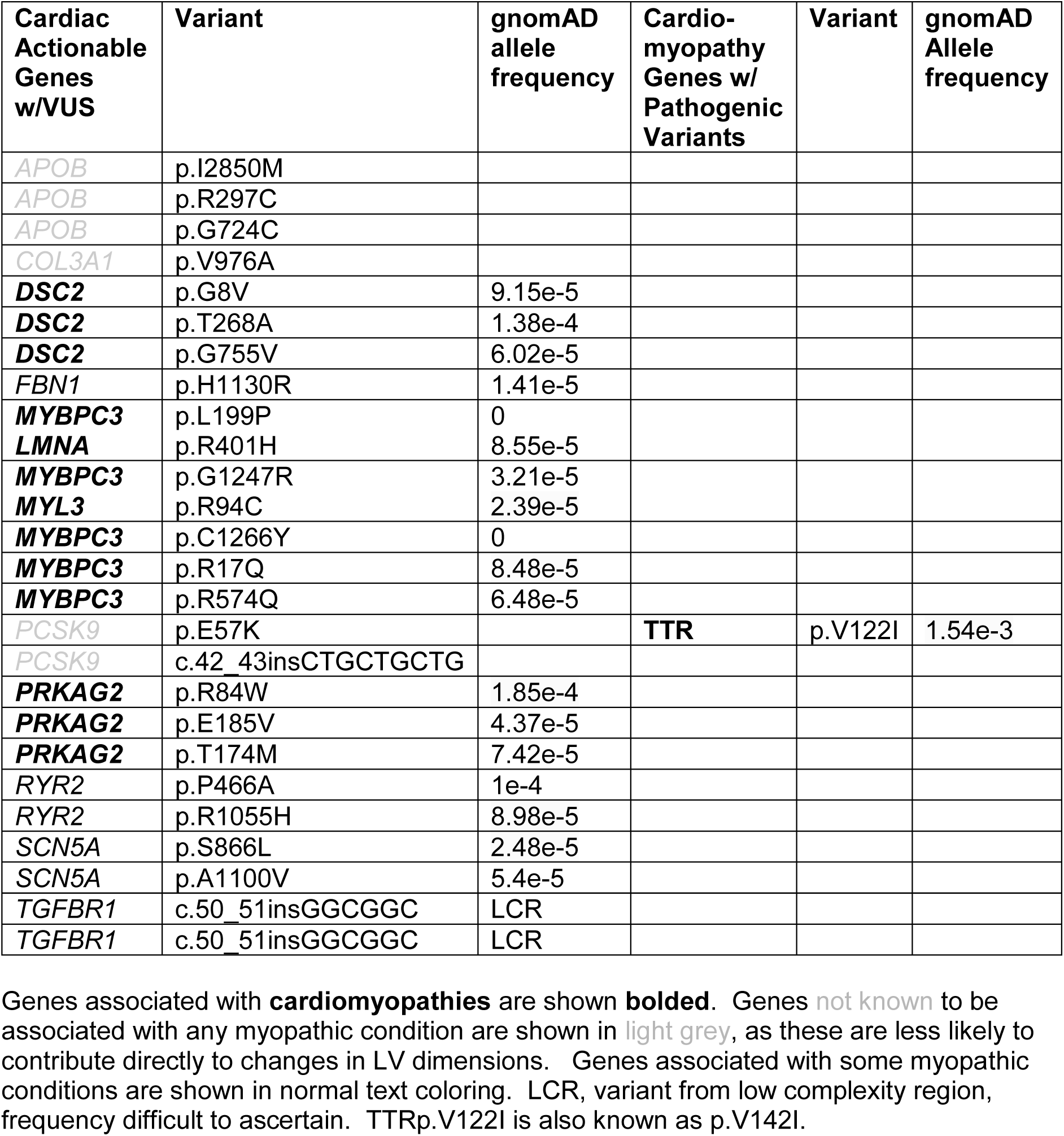
VUS found in participants with a cardiomyopathy diagnosis.

| Cardiac Actionable Genes w/VUS | Variant | gnomAD allele frequency | Cardio-myopathy Genes w/ Pathogenic Variants | Variant | gnomAD Allele frequency |
| --- | --- | --- | --- | --- | --- |
| *APOB* | p.I2850M |  |  |  |  |
| *APOB* | p.R297C |  |  |  |  |
| *APOB* | p.G724C |  |  |  |  |
| *COL3A1* | p.V976A |  |  |  |  |
| **DSC2** | p.G8V | 9.15e-5 |  |  |  |
| **DSC2** | p.T268A | 1.38e-4 |  |  |  |
| **DSC2** | p.G755V | 6.02e-5 |  |  |  |
| *FBN1* | p.H1130R | 1.41e-5 |  |  |  |
| **MYBPC3** | p.L199P | 0 |  |  |  |
| **LMNA** | p.R401H | 8.55e-5 |  |  |  |
| **MYBPC3** | p.G1247R | 3.21e-5 |  |  |  |
| **MYL3** | p.R94C | 2.39e-5 |  |  |  |
| **MYBPC3** | p.C1266Y | 0 |  |  |  |
| **MYBPC3** | p.R17Q | 8.48e-5 |  |  |  |
| **MYBPC3** | p.R574Q | 6.48e-5 |  |  |  |
| *PCSK9* | p.E57K |  | **TTR** | p.V122I | 1.54e-3 |
| *PCSK9* | c.42_43insCTGCTGCTG |  |  |  |  |
| **PRKAG2** | p.R84W | 1.85e-4 |  |  |  |
| **PRKAG2** | p.E185V | 4.37e-5 |  |  |  |
| **PRKAG2** | p.T174M | 7.42e-5 |  |  |  |
| *RYR2* | p.P466A | 1e-4 |  |  |  |
| *RYR2* | p.R1055H | 8.98e-5 |  |  |  |
| *SCN5A* | p.S866L | 2.48e-5 |  |  |  |
| *SCN5A* | p.A1100V | 5.4e-5 |  |  |  |
| *TGFBR1* | c.50_51insGGCGGC | LCR |  |  |  |
| *TGFBR1* | c.50_51insGGCGGC | LCR |  |  |  |

Genes associated with **cardiomyopathies** are shown **bolded**. Genes **not known** to be associated with any myopathic condition are shown in **light grey**, as these are less likely to contribute directly to changes in LV dimensions. Genes associated with some myopathic conditions are shown in normal text coloring. LCR, variant from low complexity region, frequency difficult to ascertain. TTRp.V122I is also known as p.V142I.

**Supplemental Table 6.** Additional Information on VUSs found in cardiomyopathy subjects.

| **Gene** | **Variant** | **Frequency** | **Diagnosis Listed in ClinVar** | **Most recent report** |
| --- | --- | --- | --- | --- |
| **DSC2** | p.G8V | 9.15e-5 | Cardiomyopathy | 2018 |
| **DSC2** | p.T268A | 1.38e-4 | Cardiomyopathy, ARVC | 2018 |
| **DSC2** | p.G755V | 6.02e-5 | ARVC | 2017 |
| **FBN1** | p.H1130R | 1.41e-5 | Marfan,TAAD | 2018 |
| **MYBPC3** | p.L199P | 0 | HCM | 2015 |
| **LMNA** | p.R401H | 8.55e-5 | CMT | 2018 |
| **MYBPC3** | p.G1247R | 3.21e-5 | Cardiomyopathy, HCM | 2018 |
| **MYL3** | p.R94C | 2.39e-5 | Cardiomyopathy | 2016 |
| **MYBPC3** | p.C1266Y | 0 | Cardiomyopathy, HCM | 2018 |
| **MYBPC3** | p.R17Q | 8.48e-5 | Cardiomyopathy, HCM | 2018 |
| **MYBPC3** | p.R574Q | 6.48e-5 | HCM, DCM, LVNC | 2018 |
| **PRKAG2** | p.R84W | 1.85e-4 | Cardiomyopathy | 2018 |
| **PRKAG2** | p.E185V | 4.37e-5 | Glycogen storage heart | 2018 |
| **PRKAG2** | p.T174M | 7.42e-5 | Glycogen storage heart | 2018 |
| **RYR2** | p.P466A | 1e-4 | Ventricular tachycardia | 2018 |
| **RYR2** | p.R1055H | 8.98e-5 | Ventricular tachycardia | 2018 |
| **SCN5A** | p.S866L | 2.48e-5 | Brugada | 2018 |
| **SCN5A** | p.A1100V | 5.4e-5 | Cardiovascular phenotype, LQTS, Brugada | 2018 |
| **TTR** | p.V122I (AKA, p.V142I) (PATH) | 1.54e-3 | Carpal Tunnel, Transthyretin Cardiomyopathy | 2018 |

ARVC, Arrhythmogenic Right Ventricular Cardiomyopathy
CMT, Charcot Marie Tooth
TAAD, Thoracic Aortic Aneurysm Dissection
LVNC, Left Ventricular Noncompaction Cardiomyopathy
MYBPC3, p.G1247R = p.G1248R

**Supplemental Figure 1.**
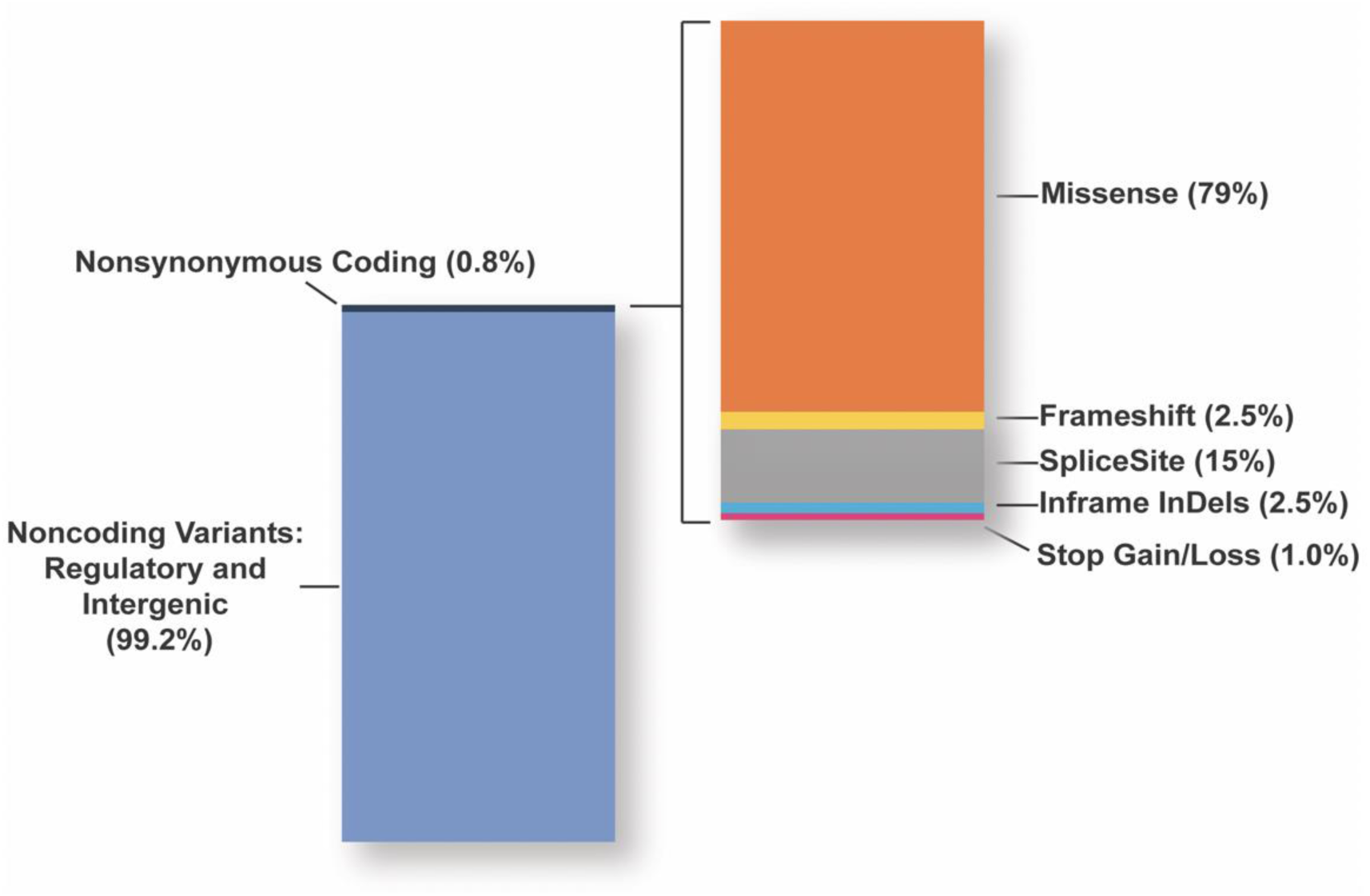
Distribution of genetic variation in the NUGene cohort as determined by whole genome sequencing. On average, 5,300,085±416,604 variants per person were identified in comparison the reference human genome. When restricting this analysis to nonsynonymous variants in the coding region, 17,282±1,234 variants per person were identified. Of these, per person there were 13,618±975 missense variants (orange), 400±31 frame shift variants (yellow), 2,500±195 splice site variants (grey); 400±35 inframe insertions or deletions (in/dels, light blue), and 200±13 variants that introduced a new stop codon or removed a stop codon (stop gain/loss, pink).

**Supplemental Figure 2.**
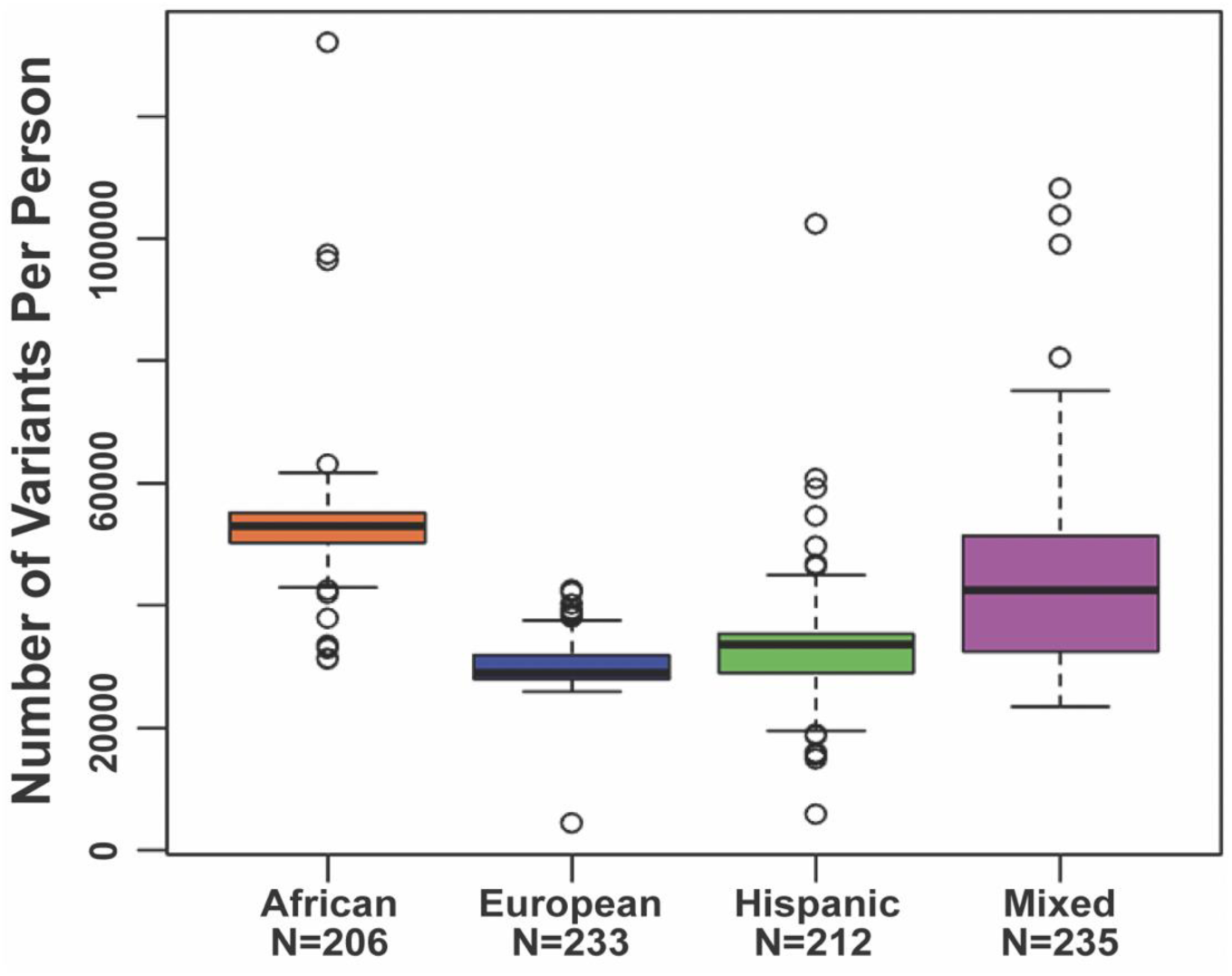
The number of variants observed only once within NUgene populations. The average number of variants that are unique to individuals in the cohort by self-reported race/ethnicity; African 52,865±8,320; European 30,217±3,992; Hispanic 32,664±7,851; and mixed race 43,513±13,199. All values are per person (p < 0.05 ANOVA across all self-reported race/ethnicities.)

**Supplemental Figure 3.**
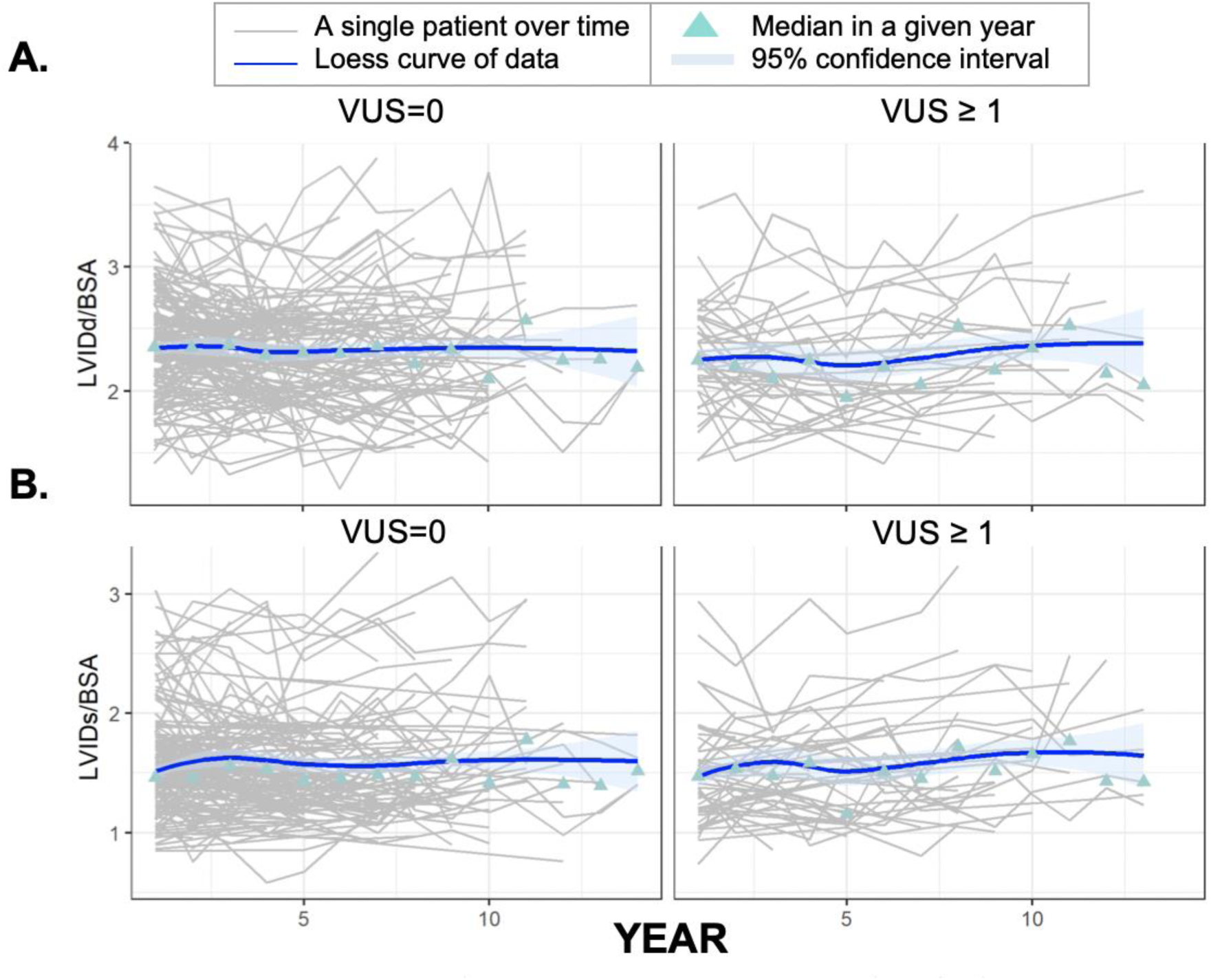
Median left ventricular dimensions correlated to VUS number in cardiac actionable genes <u>across the entire cohort.</u>. **(A)** Left ventricular internal dimension in diastole (LVIDd) corrected for body surface area (BSA) over 14 years of hospital visits by the number of variants of uncertain significance found in the cardiac actionable genes. **(B)** Left ventricular internal dimension in systole (LVIDs) corrected for BSA over 14 years of hospital visits by the number of variants of uncertain significance found in the cardiac actionable genes.

**Supplemental Figure 4.**
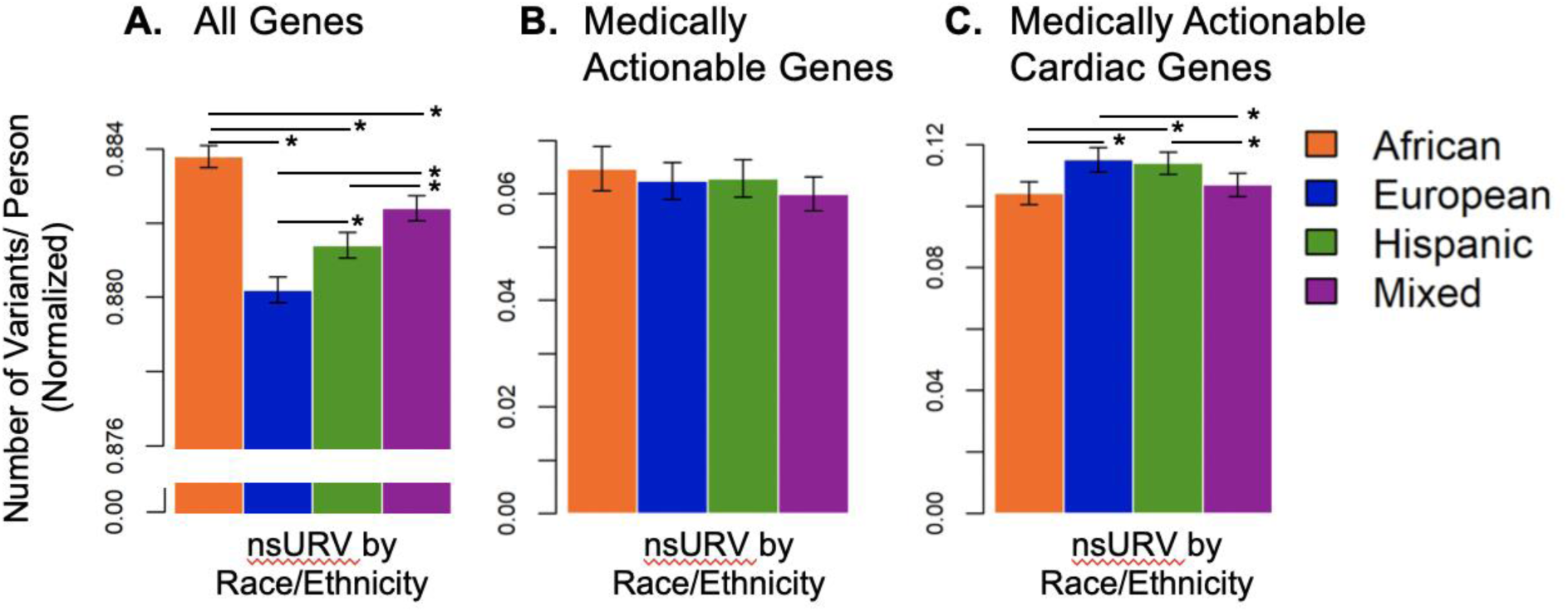
African ancestry biobank participants have significantly more unreported variants (URV). **(A)** The average number of total nonsynonymous coding variants across all genes is shown for each group based on self-reported race. Unreported variants (URV) in ClinVar were greater in African ancestry biobank participants compared to other groups (p<0.0001, ANOVA across all groups). **(B)** The average number of nonsynonymous coding variants in the 59 medically actionable genes is shown by self-reported race. The number of unreported variants (URV) in ClinVar were less in African ancestry biobank participants compared to all other groups (p<0.0001, ANOVA across all groups). **(C)** The average number of nonsynonymous coding variants in the 30 cardiac actionable genes is shown by self-reported race. The number of unreported variants (URV) in ClinVar were less in African ancestry biobank participants compared to all other groups (p<0.0001, ANOVA across all groups).

**Supplemental Figure 5.**
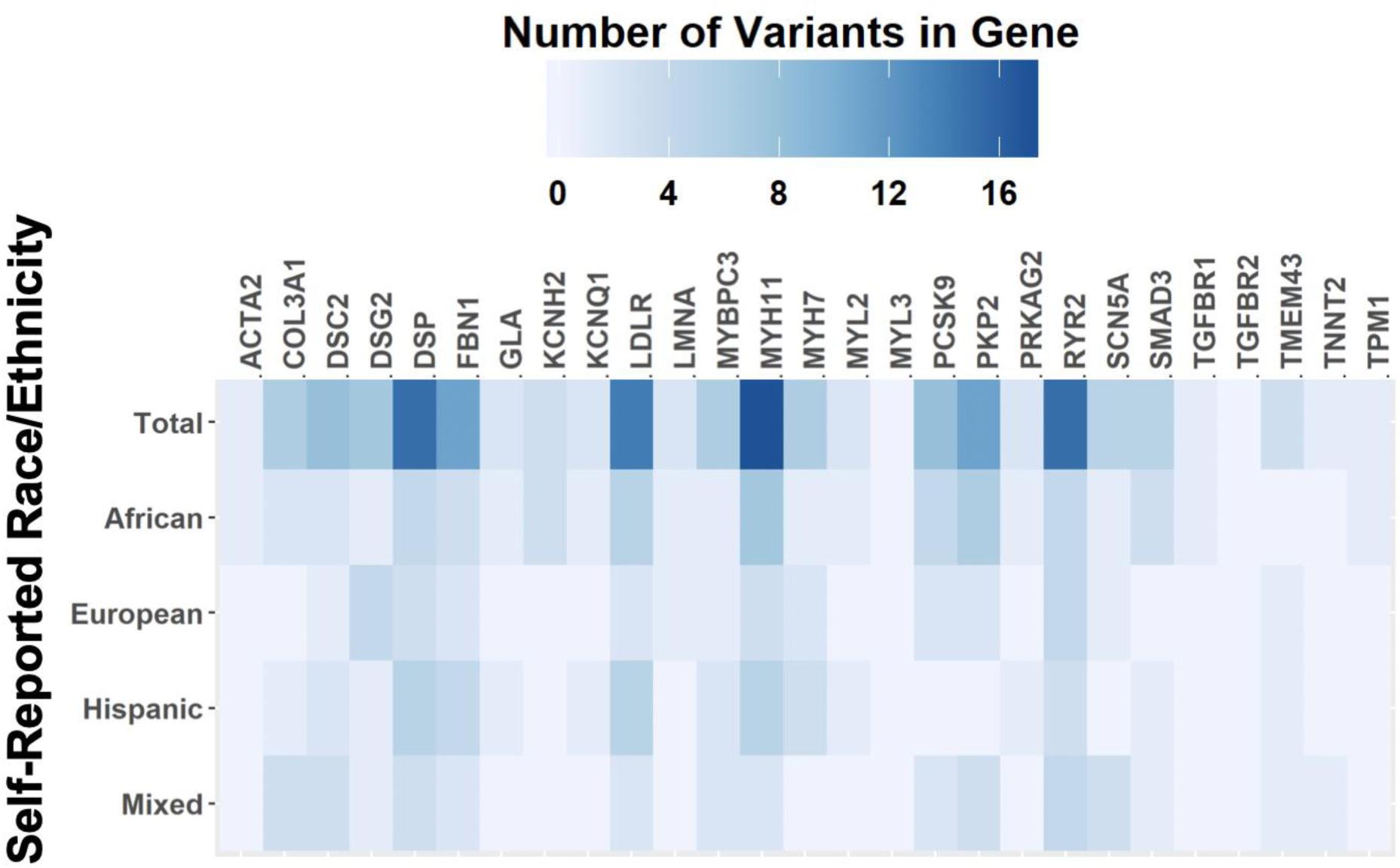
Unreported Variants in medically actionable cardiac genes by Race/Ethnicity. The number of unreported variants in cardiac actionable genes in the NUgene cohort. *APOB* is not shown.

